# R implementation of the NORMAgene data-driven qRT-PCR normalization algorithm

**DOI:** 10.64898/2026.11.15.766780

**Authors:** Grant C. O’Connell

**Affiliations:** Molecular Biomarker Core, Case Western Reserve University, Cleveland, OH, United

## Abstract

Effective data normalization is essential in producing meaningful qRT-PCR results. Traditional reference gene normalization has high risk of confounds when poorly implemented, and data-driven normalization is becoming an increasingly used and robust alternative. The NORMAgene least squares-based normalization algorithm is one of the most application flexible of existing data-driven qRT-PCR normalization approaches; however, it is currently only available in the form of an unmaintained Visual Basic for Applications (VBA) macro. In order to make this approach more widely and persistently accessible, we developed an R language implementation based on the core mathematical model and made it available as a standalone package titled ‘NORMAgene’. Here, we detail the development of this package, demonstrate it produces identical normalization results to the originally released VBA implementation, and describe how it can be used.

## Introduction

Quantitative reverse transcription PCR (qRT-PCR) remains one of the most sensitive and widely used techniques for quantification of gene expression. While qRT-PCR can produce robust results, measures can be influenced by inter-sample variations in RNA integrity, amplification efficiency, and liquid handling. As a result, normalization is required to control for such sources of experimental bias. Historically, the most commonly implemented methods of qRT-PCR normalization have been reliant on the use of reference genes [1, 2]. However, reference gene-based normalization has significant limitations; not only does it require additional measures alongside those of the target genes of interest, it can actually increase bias if poorly implemented [3–6]. As a result, there has been an increase in the use of data-driven normalization approaches as an alternative. As opposed to reference gene-based approaches wherein normalization is based solely on the observed signals of a small handful of genes which are assumed to have invariant expression, these approaches reduce artifactual technical variance via normalization factors derived by analyzing the collective pattern of signals observed across the entirety of genes measured in a given experiment [7].

Numerous benchmarking studies across a variety of applications have demonstrated that data-driven normalization approaches offer similar or superior performance to reference gene-based approaches [3, 7–10]. However, one of the reasons data-driven approaches have not become more ubiquitously used is that the most common strategies such as quantile normalization require expression data from hundreds of genes, and are thus impractical for the types of small scale experiments often performed in academic research settings. To address this limitation, Heckmann et al. developed a normalization algorithm that was specifically designed to work with smaller experimental datasets which they named NORMAgene [9]. This algorithm uses within experimental conditions least squares fits to estimate per-replicate technical variance and generate corresponding scaling factors that are ultimately applied for normalization. Like other data-driven approaches, normalization is reference gene agnostic, however, it can be carried out on datasets containing information from as few as five target genes.

Since its development, the NORMAgene algorithm has been successfully deployed in hundreds of studies across a variety of research areas [11–15], and remains one of the only data-driven qRT-PCR normalization approaches that works with small datasets. However, it is currently only available via an unmaintained Visual Basic for Applications (VBA) macro for Microsoft Excel shared alongside the original report 15 years ago. In order to make this approach more widely and persistently accessible, we sought to develop a freely available R language implementation based on the core mathematical model.

### Package build and testing

The ‘NORMAgene’ package was manually coded in the RStudio integrated development environment (Posit Software, PBC) using R version 4.5.3 (R Project for Statistical Computing) [16]. The core normalization algorithm was built using the exact mathematical equations used to build the original VBA implementation, which were retrieved from Appendix A of Heckmann et al. [9]. Documentation was written and built using markdown via the ‘roxygen2’ package [17]. A comprehensive test suite was written using the ‘testthat’ package [18], and used to verify all functions produced expected outputs and warning behavior, even in edge cases. The ‘devtools’ package [19] was used to build the final package from source and confirm it could be successfully installed and loaded in both Windows and unix-based environments.

In order to confirm that the final R package produces identical results as the original VBA implementation, we employed two real world qRT-PCR datasets previously generated by our research group. The first contains raw crossing threshold (CT) values for 12 target genes measured in total RNA isolated from 29 ischemic stroke patients and 33 neurologically normal controls, and was used in a prior benchmarking investigation comparing the NORMAgene method against traditional reference gene-based normalization [3]. The second contains raw CT values for 12 target genes measured in total RNA isolated from a single cohort of 26 ischemic stroke patients in a prior mechanistic study of the post-stroke immune response [20]. Raw CT values from the entirety of each dataset were normalized using both a copy of the original VBA macro distributed by Heckmann et al. [9] and the final R package, and the resultant outputs were tested for concordance. In order to confirm identical handling of missing values, this test was repeated after randomly removing 5% of the raw CT values from each dataset. The VBA macro was originally downloaded from https://sorensenlab.wordpress.com/2016/08/24/downloads/ in 2016 and was run using Excel version 2603 (Microsoft Inc).

### Final package, usage, and performance

The final ‘NORMAgene’ package is available for download from the Comprehensive R Archive Network (CRAN) at https://www.doi.org/10.32614/CRAN.package.NORMAgene [21]. Like the original VBA implementation, the package has the capacity to generate both normalized CT values and corresponding relative variability values, the latter of which can be used to evaluate the effect of normalization on CT variability. This metric represents the proportional change in CT value standard deviation pre to post-normalization, with values of less than one indicating a reduction in variance as a result of normalization, and values of greater than one indicating an increase in variance as a result of normalization.

The package ships three user facing functions: norma_gene(), correction_factors(), and relative_variability(). norma_gene() is directly responsible for carrying out normalization and calculating relative variability metrics. Technical replicate averaged raw CT values are provided via an input data frame appended with requisite experimental metadata including biological replicate identifiers and experimental condition information (Figure 1A). An identically structured data frame containing normalized CT values is returned, with the per-replicate correction factors applied for normalization, as well as both gene-wise and condition-wise relative variability metrics, attached as attributes. The normalized CT values can be saved as an object for downstream analysis or exported, while the associated correction factors and relative variability metrics can be accessed respectively using correction_factors() and relative variability(). Code for a typical workflow is depicted in Figure 1B, and can be run using the example raw CT values provided in Supplementary Data 1. More details regarding each individual function can be found in the manual that ships with the package.

**Figure 1.**
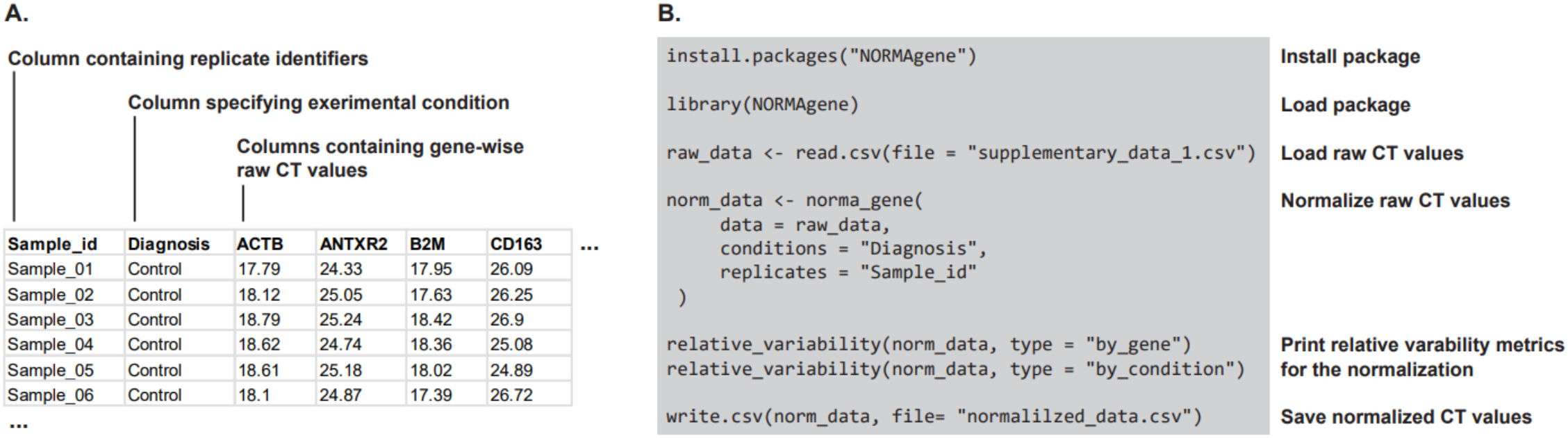
Installation and use of the ‘NORMAgene’ package. A. Input data structure. B. Example code depicting package installation and a typical normalization workflow for a gene expression profiling experiment involving multiple experimental groups.

In terms of performance, the final ‘NORMAgene’ package produced perfectly identical normalized CT values as the original VBA macro in our benchmarking tests, using both multi and single experimental condition datasets as input, including in the presence of missing values (Figure 2A). Similarly, across all tests, the final R package produced perfectly identical gene-wise and condition-wise relative variability metrics (Figure 2B). These results confirm that the R package produces a bit-by-bit computational match to the original VBA macro in terms of variable calculation, indicating that groups currently using the VBA macro can switch to the R implementation without any change in results.

**Figure 2.**
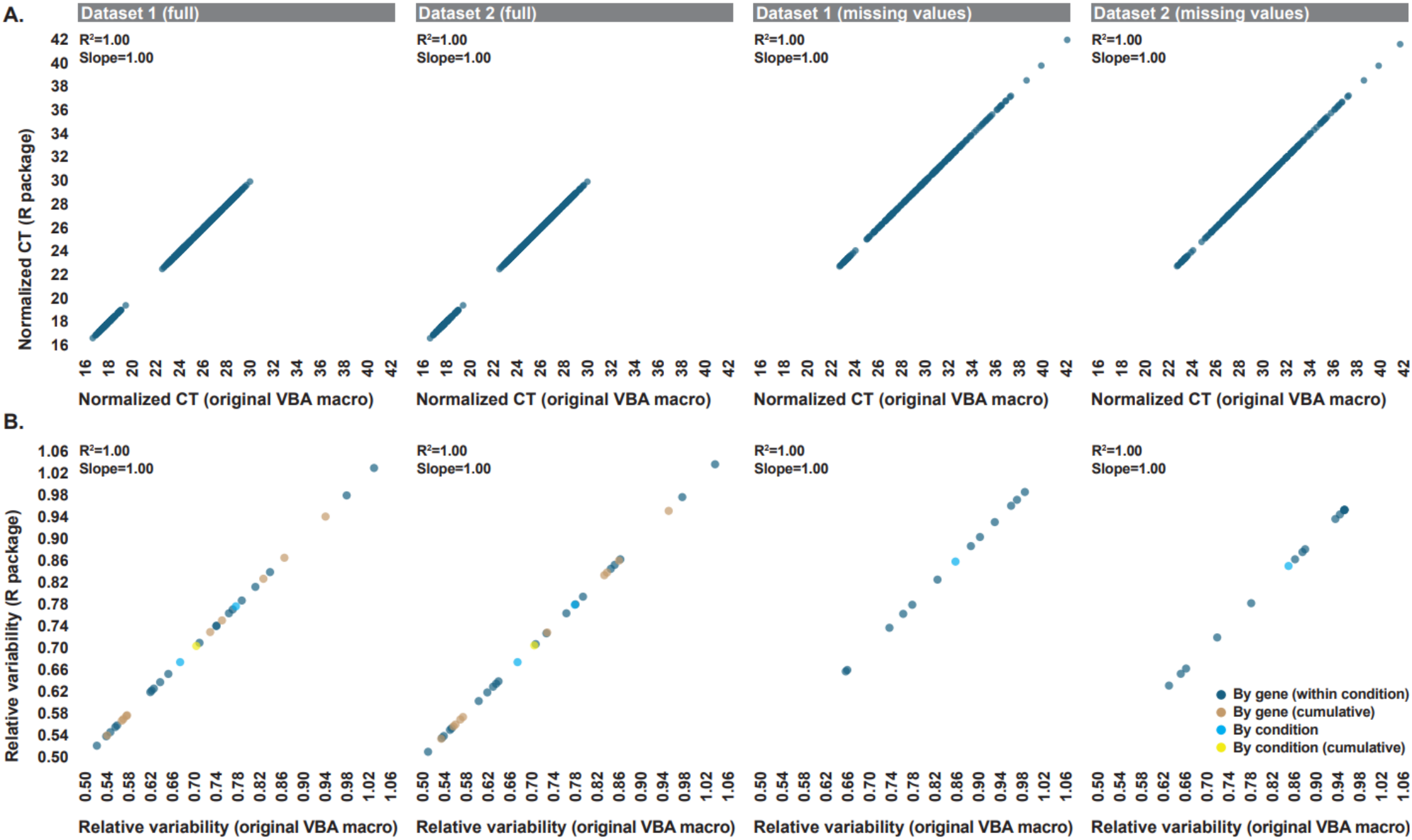
Benchmarking against the original VBA implementation. A. Correlation between normalized CT values generated by the ‘NORMAgene’ package and the original VBA implementation across both benchmarking datasets, both in the presence and absence of missing data. B. Correlation between gene-wise and condition-wise relative variability values generated by the ‘NORMAgene’ package and the original VBA implementation across both benchmarking datasets, both in the presence and absence of missing data. All correlations were assessed using Pearson’s R^2^. Slope values were generated via simple linear regression.

### Limitations

The same use-caveats associated with the original VBA implementation still apply to the ‘NORMAgene’ package, as they are associated with the underlying normalization strategy as opposed to the delivery format [3, 9]. Data from a minimum of five target genes and at least three biological replicates per experimental condition is required for reliable normalization. Given that correction factors are calculated within experimental conditions, it is imperative that group information is accurately provided. Furthermore, within condition calculation of correction factors necessitates the use of a block design with regards to upstream analytical workflow in order to facilitate accurate post-normalization inter-group comparisons of gene expression. Specimens from all experimental conditions should be processed in parallel with regards to RNA extraction, cDNA synthesis, and PCR detection; if any of these steps are carried out in batch, each batch should contain specimens from each experimental group. Similarly, there are other decisions in terms of analytical design that can maximize the amount of technical variance that is able to be detected and corrected for during normalization. For example, in large experiments, running each target gene on dedicated plates and using an identical plate position for each biological replicate across all runs allows normalization to correct for well-to-well variability in instrument signal.

## Author contributions

GOC developed the ‘NORMAgene’ package and carried out all analyses. The manuscript was written by GOC.

